# Flagellar pocket collar biogenesis: Cytoskeletal organization and novel structures in a unicellular parasite

**DOI:** 10.1101/2025.03.23.644790

**Authors:** Zelená Marie, Casas Elina, Lambert Chloé, Landrein Nicolas, Dacheux Denis, I. Abesamis Kim, Dong Gang, Varga Vladimir, Derrick. R. Robinson, Bonhivers Mélanie

## Abstract

Understanding how cells build and organize their internal structures is a fundamental question in biology, with important implications for human health and disease. Trypanosomes are single-celled flagellated parasites that cause life-threatening diseases in human and animals. Their survival relies on a specialized compartment called the flagellar pocket (FP), which serves as a gateway for nutrient uptake, and immune evasion. The formation and function of the FP are supported by an intricate cytoskeletal structure known as the flagellar pocket collar (FPC). However, the mechanisms underlying its assembly remain poorly understood.

In this study, we used cutting-edge ultrastructure expansion microscopy (U-ExM) to investigate FPC biogenesis in *Trypanosoma brucei*. We mapped the formation of the new microtubule quartet (nMtQ) alongside flagellum growth, providing new insights into its assembly. Additionally, we tracked the localization dynamics of key structural proteins - BILBO1, MORN1, and BILBO2 - during the biogenesis of the FPC and the hook complex (HC). Notably, we identified two previously undetected structures: the proFPC and the transient FPC-interconnecting fibre (FPC-IF), both of which appear to play crucial roles in linking and organizing cellular components during cell division.

By uncovering these novel aspects of FPC biogenesis, our study significantly advances the understanding of cytoskeletal organization in trypanosomes and opens new avenues for exploring the functional significance of these structures.

## Introduction

The biogenesis of cytoskeletal structures is a fundamental process that shapes eukaryotic cell architecture and function. It is essential for maintaining cell shape, facilitating intracellular transport, driving cell division, and enabling cellular motility. This process involves the synthesis of monomeric proteins, their polymerization into higher-order assemblies, and the coordinated action of various regulatory proteins and signalling pathways. In many protist parasites, which possess highly organized cytoskeletal architectures, these mechanisms are particularly critical as they transition between hosts, undergoing distinct morphological stages. Trypanosomatids are flagellated protist parasites responsible for devastating human and animal diseases, including human African sleeping sickness, Nagana, Chagas disease, and leishmaniasis. Studies on trypanosome morphology and biochemistry have revealed a dense subpellicular microtubule cytoskeleton (1), which forms a corset-like structure, restricting endo- and exocytosis. To circumvent this limitation, trypanosomes possess a specialized invagination of the plasma membrane at the base of the flagellum, known as the flagellar pocket (FP). The FP is essential for nutrient uptake, protein secretion, immune evasion, and cell morphogenesis (2–4). Similar pocket-like structures exist at the base of cilia in other organisms, including mammalian primary and motile cilia (5).

The FP is a conserved feature among trypanosomatids, and these parasites have evolved specialized structures within the FP distal region to enhance its function. This region, referred to as the neck, is tightly enclosed around the flagellum by the flagellar pocket collar (FPC), a key cytoskeletal structure (6–8). The FP and FPC are closely associated with the Hook Complex (HC), a cryptic MORN1 protein containing structure (9) and a distinct set of four microtubules known as the microtubule quartet (MtQ) (Sherwin and Gull, 1989; Vickerman, 1969).

In G1-phase cells, the MtQ arises between the mature and pro-basal bodies of the flagellum, wraps around the FP, and extends along the entire length of the cell body parallel to the flagellum. Although its precise functions remain unclear, the MtQ is believed to help define FP architecture and assist in the formation of the new FP during cell division.

The cytoskeleton-associated HC, located distal and adjacent to the FPC, forms a hook-like structure in close proximity to the FPC (reviewed in (9)). The first identified HC protein, MORN1, is essential in the bloodstream form of *Trypanosoma brucei*, and its depletion disrupts the endomembrane system, although its molecular function remains unclear (10,11).

BILBO1, a kinetoplastid-specific protein, was the first FPC protein to be identified (6). It is a protein that polymerizes both *in vivo* and *in vitro* (12,13) serving as a the structural backbone of the FPC and interacting with multiple binding partners including TbKINX1b (a basal body kinesin), BILBO2 (an FPC protein), and FPC4 (a MtQ-binding protein) (14–16). These interactions suggest a strong functional link between BILBO1 and microtubule-based structures such as the MtQ. Notably, BILBO1 also interacts with BHALIN, a protein localized to the HC, reinforcing the tight structural association between the FPC and the HC (17). To date, BILBO1 remains the only protein proven to be essential for the overall biogenesis of the FPC, FP, and HC, positioning it as a master organizer for these structures (6,16,17).

Tomography studies have characterized the morphology of the FPC and its association with cytoskeletal structures, including the MtQ, highlighting their close association (18–20). Importantly, FP formation begins with the emergence of a ridge between the old and new FPs (20), but its biogenesis is dependent on the FPC (Bonhivers et al., 2008). However, the precise mechanisms governing FPC formation during the cell cycle remain largely unknown.

In this study, we characterize the cell cycle-related stages of FPC formation at high resolution using ultrastructure expansion microscopy (U-ExM), revealing that FPC biogenesis occurs *de novo* yet remains closely associated with the existing FPC and MtQ. Additionally, we identify a previously uncharacterized transient filamentous structure, termed the FPC-interconnecting fibre, which contains several FPC-related proteins. Finally, we explore the role of FPC biogenesis in FP formation and function, providing new insights into the assembly and regulation of these essential cytoskeletal structures.

## Results

### Spef1 tracking and U-ExM reveal sequential MtQ formation and flagellum elongation during the cell cycle

To gain a clearer understanding of MtQ formation, we leveraged the advanced imaging capabilities of ultrastructure expansion microscopy (U-ExM) in *T. brucei* (15,21,22) and combined this approach with the generation of specific cell lines. We first developed a cell line expressing an endogenously tagged _HA_Spef1 protein, which labels the MtQ between the basal bodies and the FPC as previously described (23). U-ExM was then applied to detergent-extracted _HA_Spef1-expresssing cells (cytoskeletons). Co-immunolabelling of Spef1 and tubulin allowed us to visualise the formation of the new MtQ (nMtQ) and track new flagellum (nFg) elongation throughout the cell cycle (Figure 1).

**Figure 1.**
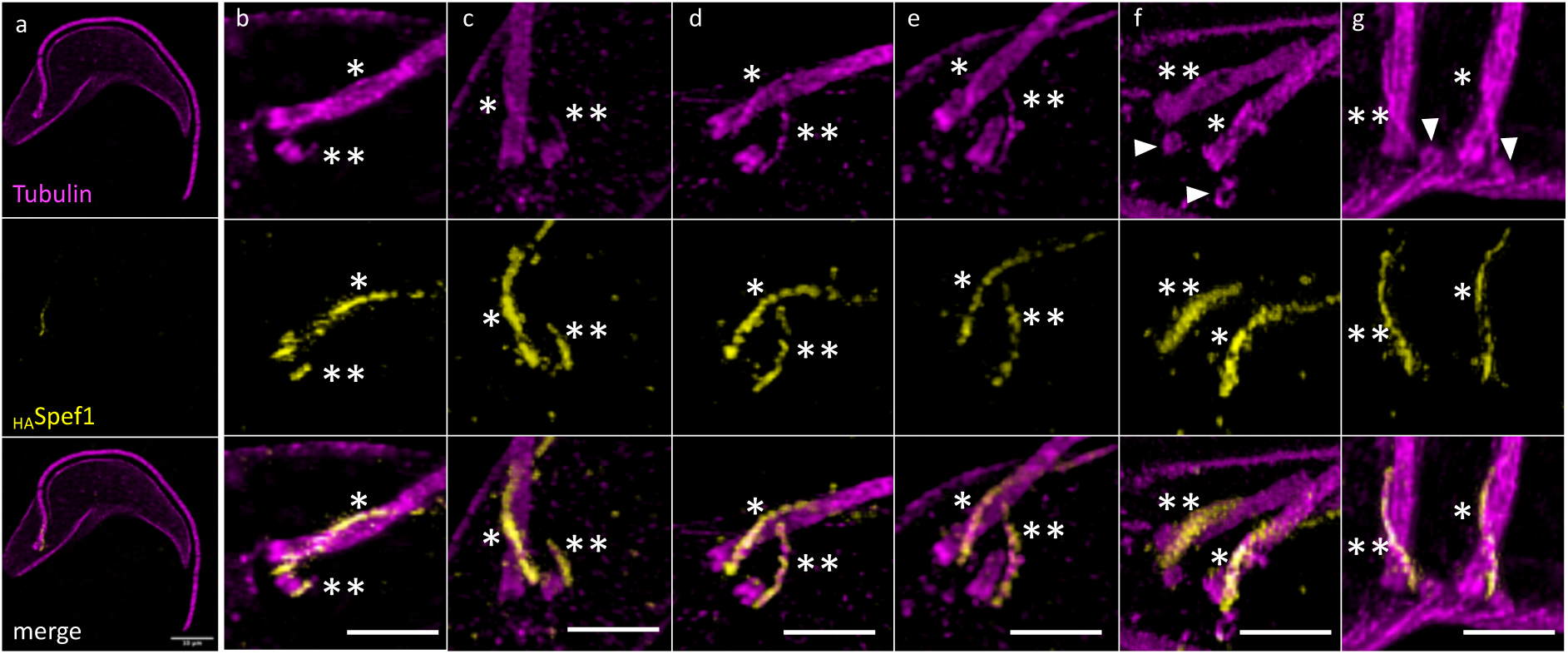
U-ExM images showing co-immunolabelling of tubulin and _HA_Spef1 on expanded cytoskeletons. Tubulin (magenta), _10HA_Spef1 (yellow) (snapshots from 3D viewer). *: old MtQ. **: new MtQ. Arrowheads: new pro-basal bodies. Scale bars: 10μm (a), 5μm (b-g).

In G1-phase cells, Spef1 was detected on the old MtQ (oMtQ) (Figure 1a). As the cell cycle progressed, Spef1 labelling appeared on the newly forming nMtQ (Figure 1b). This nascent nMtQ was not located within the inter-basal body region but instead positioned adjacent to and on the outer periphery of the pro-basal body (proBB). As the nMtQ extended, it remained continuously labelled by Spef1 and gradually approached the oMtQ (Figure 1c-e).

Consistent with previous observations (20), the nFg subsequently rotates around the old flagellum (oFg) while remaining attached to it, repositioning the nMtQ to a more posterior location within the cell (Figure 1f-g). Following this rotation, two new proBBs were clearly observed: one associated with the old basal body and the other with the new basal body. Notably, the nMtQ extended to the oMtQ before the formation of the new transition zone (Figure 1d), indicating that nMtQ formation and extension precede new flagellum elongation.

### Sequential BILBO1 localization reveals key steps in FPC biogenesis and identifies novel proFPC and FPC-interconnecting structures

Previous electron microscopy and epifluorescence studies using a polyclonal antibody against BILBO1 have localized this protein to the FPC, as well as along the MtQ and basal bodies (6,14,10). To gain further insights into FPC formation, we used U-ExM to examine BILBO1 localization throughout the cell cycle in relation to Spef1 and tubulin labelling (Figure 2).

**Figure 2.**
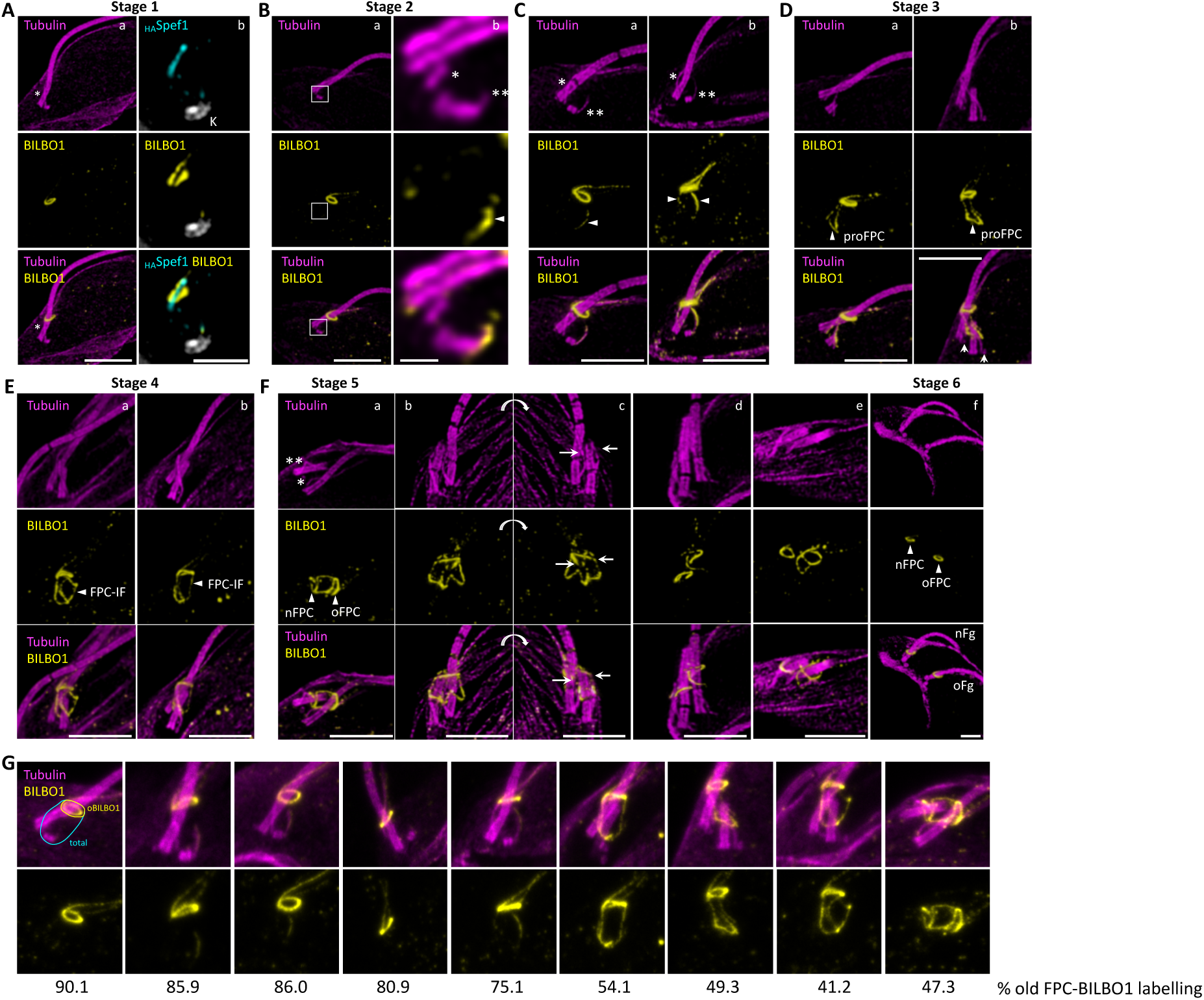
U-ExM immunofluorescence images showing localization of BILBO1 localization in detergent-extracted *T. brucei* cells. **A. a.** Immunolabeling of tubulin (magenta) and BILBO1 (yellow) labelling. BILBO1 localizes at the FPC only. * indicates the MtQ. **b.** Co-labeling of BILBO1 (yellow) and _HA_Spef1 labelling (cyan). A single plane from a Z-stack of Spef1 (_HA_Spef1) and BILBO1 co-labelling reveals that BILBO1 forms a flat corkscrew structure, with Spef1 (and thus the MtQ) threading through the corkscrew. The kinetoplast was labelled with Hoechst (K, grey). **B-F.** Co-labeling of tubulin (magenta) and BILBO1 (yellow) at different cell cycle stages. **B. a.** The boxed region is enlarged in b. **b.** * represents the old MtQ, and ** represents the new growing MtQ. The arrowhead indicates the BILBO1-positive labelling at the new MtQ. **C.** Arrowheads indicate the BILBO1-positive labeling on both the nMtQ and oMtQ. The old MtQ (*) and new MtQ (**) are indicated. **D.** In a, arrowheads indicate the tubulin-negative pro-FPC structure. In b, arrows indicate the new formed BBs. **E.** Arrowheads denote the FPC-interconnecting fibre (FPC-IF). **F.** Arrowheads indicate the new and old FPCs. nFg: new flagellum. oFg: old flagellum. In b-c, arrows indicate the extremities of the oFPC. **G.** Quantification of BILBO1 fluorescence intensity in representative images, comparing the old FPC (yellow area) to total BILBO1 labelling (cyan area). Scale bars: 5 μm.

In early G1-phase cells (Stage 1, Figure 2 Aa), BILBO1 appears as a flattened corkscrew or spring washer-like structure at the FPC. The MtQ, labelled with Spef1, threads through the groove of the FPC (Figure 2 Ab). In addition to the primary FPC-associated signal, weaker BILBO1 labelling is observed distally from the FPC in two linear structures: one along the MtQ (Figure 2) and another corresponding to the _myc_CAAP1-decorated centrin arm (24) (Figure S1). As soon as the nMtQ becomes detectable, BILBO1 colocalizes with it, adjacent to the proBB (Stage 2, Figure 2Ba, b). Notably, tubulin and BILBO1 co-labelling confirms that the FPC remains tubulin-negative throughout the cell cycle, indicating that tubulin is not a component of the FPC or involved in its biogenesis. Interestingly, at this stage, BILBO1 is absent from the oMtQ. However, as the nMtQ extends towards the old FPC, it remains BILBO1-positive (Figure 2Ca). Once the nMtQ reaches close proximity to the old FPC, a BILBO1 signal is also observed on the old MtQ (Figure 2Cb, c). The absence of cells in an intermediate state suggests that BILBO1 recruitment to the oMtQ occurs rapidly.

As the nFg grows, a newly-formed BILBO1-positive structure emerges, wrapping around the region of the transition zone of the growing flagellum and forming a semi-circular structure that connects the old and new MtQs (Stage 3, Figure 2D). This structure, which we termed the proFPC, serves as the foundation for the formation of the new FPC assembly. At this stage, tubulin labelling reveals two newly formed proBBs (Figure 2Db, arrows).

As the nFg continues to grow and approaches the oFPC, a novel BILBO1-positive, tubulin-negative fibrous structure emerges, linking the old and new FPCs (Stage 4, Figure 2E). This structure, which we termed the FPC-interconnecting fibre (FPC-IF), represents a previously uncharacterized component of FPC biogenesis. The precise mechanism by which BILBO1 is targeted to these structures remains unclear.

As the nFg rotates around the old flagellum before exiting the FP, its distal tip remains attached to the side of the old flagellum *via* the flagellum connector (FC) (20,25). This rotation, coupled with the movement of the nBB repositions the nMtQ to a more posterior location with the cell. Consequently, the nFPC adopts its characteristic corkscrew-like morphology (Stage 5, Figure 2F). Notably, at this stage, the oFPC now adopts an open-circle configuration, creating an enlarged gap (Figure 2Fb-c). This gap likely serves as the passage of the growing nFg as it exits the oFPC while remaining attached to the old flagellum *via* the FC.

Following BB segregation and subsequent separation of the nFg from the oFg, we observed the disappearance of MtQ-associated BILBO1 labelling and the loss of the FPC-IF. This suggests two key events: (i) cessation of BILBO1 targeting, recruitment or transport (if any) to the FPCs, and (ii) the transient nature of the FPC-IF (Stage 6).

To determine whether the quantity of BILBO1 at the oFPC remains stable throughout the cell cycle, we quantified its fluorescence intensity in two distinct regions: one directly surrounding the oFPC (yellow area) and another encompassing the oFPC and extending towards the BB region (cyan area) (Figure 2G). The total fluorescence intensity (set at 100%) includes signal from the oFPC, the FPC-IF and the MtQ structures. With the exception of the G1-phase cells (where no nFPC is being formed) and the period when the FPC-IF is detected, the oFPC consistently accounts for approximately 40% of the total BILBO1 fluorescence, remaining close to 50% throughout the cell cycle. This indirectly suggests that nFPC biogenesis occurs *de novo*, rather than through redistribution of BILBO1 from the oFPC.

### Cytoskeletal dynamics and FPC-IF formation: shaping the Flagellar Pocket

To investigate the biogenesis of the FP and localize the proFPC and the FPC-IF relative to the FP membrane, we performed immunolabelling with a polyclonal antibody against BILBO1 (10) on expanded whole cells (Figure 3). This is complemented with NHS-Ester Atto 594 labelling of cellular proteins (26) which delineated the FP (purple, Figure 3, Figure S2c) as an empty space (Figure 3, Figure S2e-g). This method successfully revealed the localization of key FP-associated cytoskeletal structures, including the flagellum, basal bodies, the MtQ, the FPC, and FP boundaries (Figure S2). We mapped each step of FPC biogenesis, including the formation of the nMtQ (Figure 3Ab), proFPC (Figure 3Ac-e), new flagellum (Figure 3Ac-i), FPC-IF (Figure 3Ad, e), and nFPC (Figure 3A, g-i), in relation to the FP.

**Figure 3.**
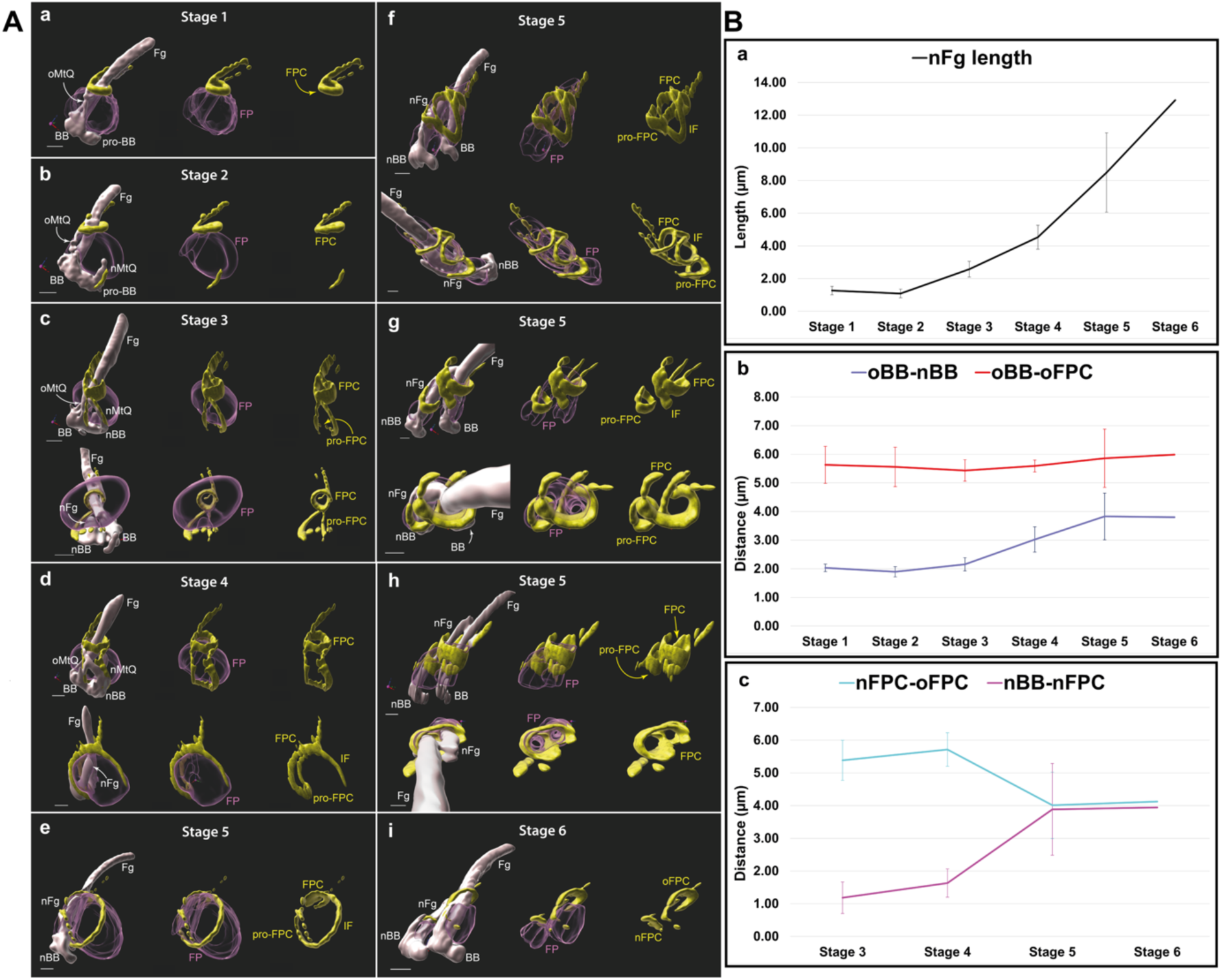
A. Representation of the flagellar pocket region in successive stages of flagellar pocket division and new flagellar pocket collar formation. These models were segmented from whole, expanded cells labelled with fluorescent NHS ester (Atto 594) and BILBO1 rabbit polyclonal antibody, followed by an anti-rabbit antibody (Alexa 488). Microtubule-based structures (grey) were segmented from NHS ester data by manual masking and thresholding. **Scale bar**: 2 µm physical distance (correspond to 0.43 µm after correction for the 4.6 fold expansion factor). **Fg** – proximal region of the old (mature) flagellum, **nFg** – new (growing) flagellum, **BB** – mature basal body belonging to the old flagellum, **pro-BB** – probasal body, **nBB** – new mature basal body, **oMtQ** – old microtubule quartet, **nMtQ** – new microtubule quartet. The flagellar pocket (FP, purple) was segmented manually from NHS ester data. BILBO1 antibody signal (yellow) is represented by intensity thresholding. **FPC** – flagellar pocket collar, **pro-FPC** – newly forming flagellar pocket collar, **oFPC** – old flagellar pocket collar, **nFPC** – new flagellar pocket collar, **IF** – interconnecting fibre. Progression of FP division: **a** G1-phase cell: A single flagellum is present before the formation of the nMtQ and nFg. BILBO1 (yellow) forms a loop around the Fg (FPC) and follows a part of the oMtQ distally from the FPC. **b** Early MtQ formation: Cells with a single flagellum cell were used to collect images of growing nMtQ and show that BILBO1 (yellow) is recruited alongside the growing nMtQ. **c.** nMtQ reaches the FPC: A nFg starts to assemble, and BILBO1 signal (yellow) is detectable on both the oMtQ and the forming pro-FPC at the base of the nFg. **d** nFg protrusion: A cell with nFg protruding into the FP, but not yet reaching the FPC. BILBO1 (yellow) forms and interconnecting fibre (IF), which originates from the FPC and follows the surface of the FP. **e** IF connection to proFPC: A later stage of d, the IF connects to the pro-FPC. **f** nFg rotation: The growing nFg rotates counter-clockwise around the oFg when viewed from the basal body towards the flagellum. The nBB is now positioned posterior to the BB. The FP starts to divide into two compartments, and the pro-FPC moves distally away from the nBB. **g** nFg approaches the FPC: The FPC loop relaxes to accommodate the nFg. **h** nFg reaches the FPC: At this stage, both the Fg and the nFg are enclosed within the FPC. However, in later stages, the nFg is only surrounded by BILBO1 at the nFPC. **i** nFg extends past the FPC: The nFPC is now fully formed and separates from the original FPC, which is now designated as of oFPC. **B. Graphs representing measurements of the data presented in A.** The measured distances shown in the graphs are uncorrected by the isotropic ∼ 4.6-fold expansion factor. Each plotted value represents the average of all values measured within each stage ± standard deviation. For the purpose of the measurements, the stages of nFg growth and FP division were divided into six stages: **Stage 1**: cells before nMtQ formation (represented by 3 Aa), **Stage 2**: cells with a forming nMtQ, but not yet forming nFg (represented by 3Ab), **Stage 3**: cells with nMtQ reaching the FPC and the nFg starting to form (represented by 3Ac), **Stage 4**: cells with nFg with a fully formed transition zone, but not yet reaching the FPC (represented by 3Ad), **Stage 5**: cells with nFg about to reach or reaching the FPC (represented by 3Ae-h), **Stage 6**: cells with nFg past the FPC and fully divided FP before the two FPs move apart (represented by Ai). The following number of cells were measured: Stage 1: 3 cells, Stage 2: 3 cells, Stage 3: 3 cells, Stage 4: 3 cells, Stage 5: 6 cells, Stage 6: 1 cell. **a** – The progress of new flagellum (nFg) growth throughout measured stages. **b – Dark blue line:** distances between the new basal body (nBB, in some stages pro-BB) and the oBB. The distance was measured from centre to centre from the most distal of each basal body. The distance increases in stages 4-6, suggesting the basal bodies start moving apart in stage 4. **Red line:** distances between the old basal body (oBB) and the old FPC (oFPC). The distance between the oBB and the nFPC was measured at the furthest point of the nFPC from the oBB. The measurements suggest that the distance between the oBB and oFPC remains constant. **c** – **Cyan line:** Distances between the nFPC (or pro-FPC) and the oFPC. The distances were measured between the furthest point of the oFPC relative to the oBB and the apex of the pro-FPC in stages 3 and 4, and between the furthest point of the oFPC relative to the oBB and the furthest point of the nFPC relative to the nBB in stages 5 and 6. The distance between both FPCs decreases slightly between stages 4 and 5. **Magenta line:** The distance between the nFPC (pro-FPC) and the nBB. The distance between the nFPC and nBB is increased between stages 3 and 5.

Whilst the precise timing of FPC-IF formation remains uncertain, our imaging confirmed that the proFPC forms in the cytosolic compartment beneath the base of the FP, and connects the two MtQ structures. Consistent with the previous U-ExM cytoskeletal data (Figure 2), the FPC-IF extends along the anterior side of the FP connecting the proFPC and the oFPC. Following nFg rotation, the FPC-IF disappears (Figure 3A, h), coinciding with a corresponding loss of most BILBO1 labelling on the MtQs (Figure 3A, g). Notably, we observed the relaxation of the oFPC into an open conformation (Figure 3A, i), with this structural transition facilitating nFg exit from the old FPC despite remaining attached to the oFg *via* the FC. Additionally, two distinct FPs were observed as soon as the nFPC was formed (Figure 3Ag).

Measurements in whole cell samples showed that the new flagellum elongates from stage 2 in a linear fashion (Figure 3Ba) and the distances between the oBB and the oFPC indicate that the mature FPC remains relatively stationary along the oFg during the cell cycle (Figure 3Bb). As previously described in (27), the oBB-nBB distance increases during the cell cycle (Figure 3Bb). In contrast, the distance between the nBB and the proFPC/nFPC reveals that the proFPC forms early in the process of nFg growth (Figure 3B a and c). The distance between the nBB and nFPC increases between stage 4 and stage 5, and coincides with the timing of nFg rotation, before stabilizing between stages 5 and 6 (Figure 3Bc). These observations suggest three possible scenarios: (i) the proFPC moves upwards along the growing nFg, (ii) the nBB moves downward into the cell. However, it is most likely that both processes occur simultaneously or at specific time points. Indeed, the first scenario is supported by the observed reduction in the distance between the nFPC and oFPC from stage 4 to stage 5, suggesting that the nFPC progressively moves closer to the cell surface (Fig. 3Bd). The second scenario is supported by the relatively large change in the nBB-nFPC distance between stage 4 and 5 (Figure 3Bc). Additionally, these observations indicate that the new flagellar pocket is not fully distinguishable morphologically until the nFPC is completely formed (Figure 3Af), implying that pocket completion depends directly on the formation of the nFPC.

### MORN1 localization during the cell cycle reveals parallel biogenesis pathways for the FPC and the HC

In G1-phase expanded cytoskeletons, MORN1 localized to the bilobe structure (9), which comprises the HC and the centrin arm, positioned adjacent to the distal side of the FPC (Figure 4Aa). This localization pattern is consistent with previous findings (10,15). While technical limitations prevented direct co-labelling of MORN1 and BILBO1, the MORN1 localization pattern closely mirrored that of BILBO1 throughout the cell cycle. In addition to marking the old HC (oHC), MORN1 was detected in detergent-extracted cells along both MtQs (Figures 4Ab and c), around the newly forming axoneme (proFPC Stage 3, Figure 4Ad), and later within a filamentous structure connecting to the oHC (FPC-IF Stage 4, Figure 4Ae). This sequential localization pattern culminated in the eventual separation of the old and new HC structures (Figure 4Af). The striking similarity between MORN1 and BILBO1 labelling strongly suggests that MORN1 is consistently present within BILBO1-positive structures throughout the cell cycle, highlighting the functional interplay between the two structures.

MORN1 labelling in expanded whole cells (Figure 4B) further emphasized the similarities between FPC and the HC formation, showing MORN1-decorated MtQs and the emergence of a proFPC-like structure (named proHC in Figure 4Be) linking the two MtQs. This structure subsequently migrates away from the nBB, giving rise to an nHC. However, in these expanded whole cells, MORN1 was not detected on the nMtQ prior to its presence on the oMtQ, nor was it observed on the FPC-IF structure. This discrepancy is most likely due to differences in labelling sensitivity, as MORN1 is clearly detected on these structures in detergent-extracted cells (Figure 4A).

**Figure 4.**
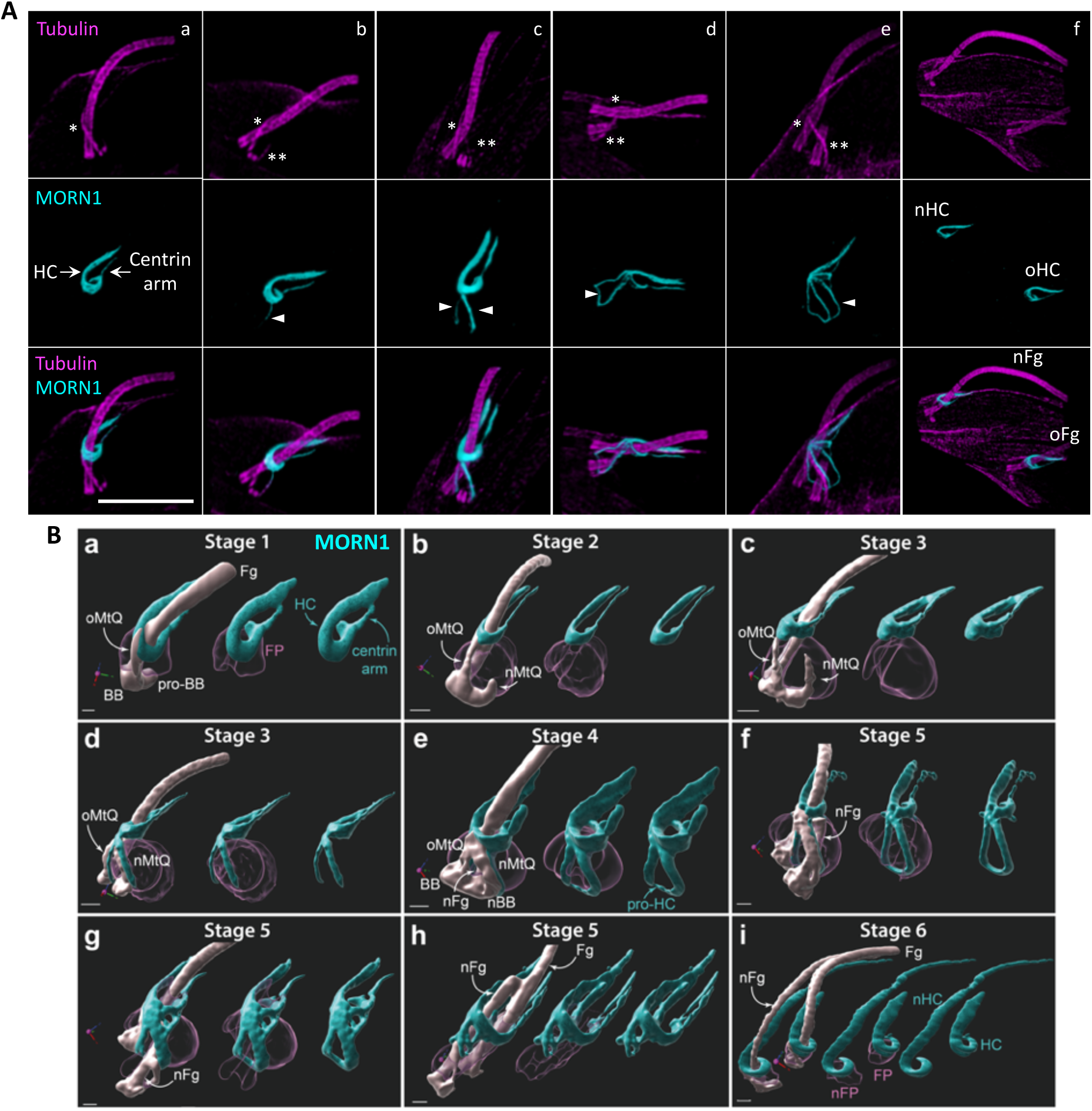
U-ExM immunofluorescence images depicting the formation of the Hook Complex (HC). **A.** Localization of MORN1 in detergent-extracted *T. brucei* cells. Expanded cytoskeletons were labelled with tubulin (magenta) and MORN1 (cyan). (a) The HC and the centrin arm of the bilobe structure are indicated by arrows. The old MtQ is marked with a single asterisk. (b) The new growing MtQ is marked with two asterisks, with MORN1 detected exclusively on the growing new MtQ (arrowhead). (c) MORN1 is detected at both the new and the old MtQ (arrowheads). (d) The pro-FPC structure is MORN1 positive (arrowhead). (e) The FPC-interconnecting fibre is also MORN1 positive (arrowhead). (f) At the final stage of the cell cycle, the two flagella separate with the HC and centrin arm. Scale bar: 5 μm. **B.** Segmented views of the flagellar pocket region at sequential stages of flagellar pocket division and HC formation in expanded whole cells labelled with anti-MORN1 and NHS ester. The oMtQ is marked with a single asterisk, while the new growing MtQ is marked with two asterisks. The arrowheads indicate the MORN1 labelling on the MtQs (d) and the proFPC-like structure (e). Scale bars: 2 μm. **Scale bar:** 2 µm physical size (corresponding to 0.43 µm after correction for the ∼ 4.6-fold expansion factor). **Fg** – proximal region of the old (mature) flagellum, **nFg** – new (growing) flagellum, **BB** – mature basal body belonging to the old flagellum, **pro-BB** – probasal body, **nBB** – new mature basal body, **oMtQ** – old microtubule quartet, **nMtQ** – new microtubule quartet. The flagellar pocket (**FP**, purple) was segmented manually from NHS ester data. MORN1 antibody signal (cyan) is represented by intensity thresholding. **HC** – hook complex, **pro-HC** – newly forming hook complex, **nFP** – new flagellar pocket.

Notably, MORN1 was never observed at the BB-proximal end of the MtQ in either detergent-extracted cytoskeleton or whole-cell samples. As previously described, MORN1 appeared only when the nMtQ reached the oFPC level, indicating that BILBO1 associates with the MtQs prior to MORN1. Interestingly, Figure 4B panel G clearly shows the formation of a distinct, flask-shaped nFP forming as the distance between the nBB and the proFPC increases. This observation reinforces the idea that the FPC not only is required to maintain the pocket neck in a closed state but also actively contributes to FP formation.

### BILBO1 and BILBO2 colocalization reveals the distinct molecular composition of the proFPC and FPC-IF

In this study, we identified three previously uncharacterized cytoskeletal structures, the proFPC, the pro-HC and the FPC-IF. Our analysis revealed that they are tubulin-negative but contain BILBO1 and most likely MORN1. Furthermore, the absence of a Spef1 signal within these structures indicates that they lack this known MtQ-associated protein (Figure 5A).

**Figure 5.**
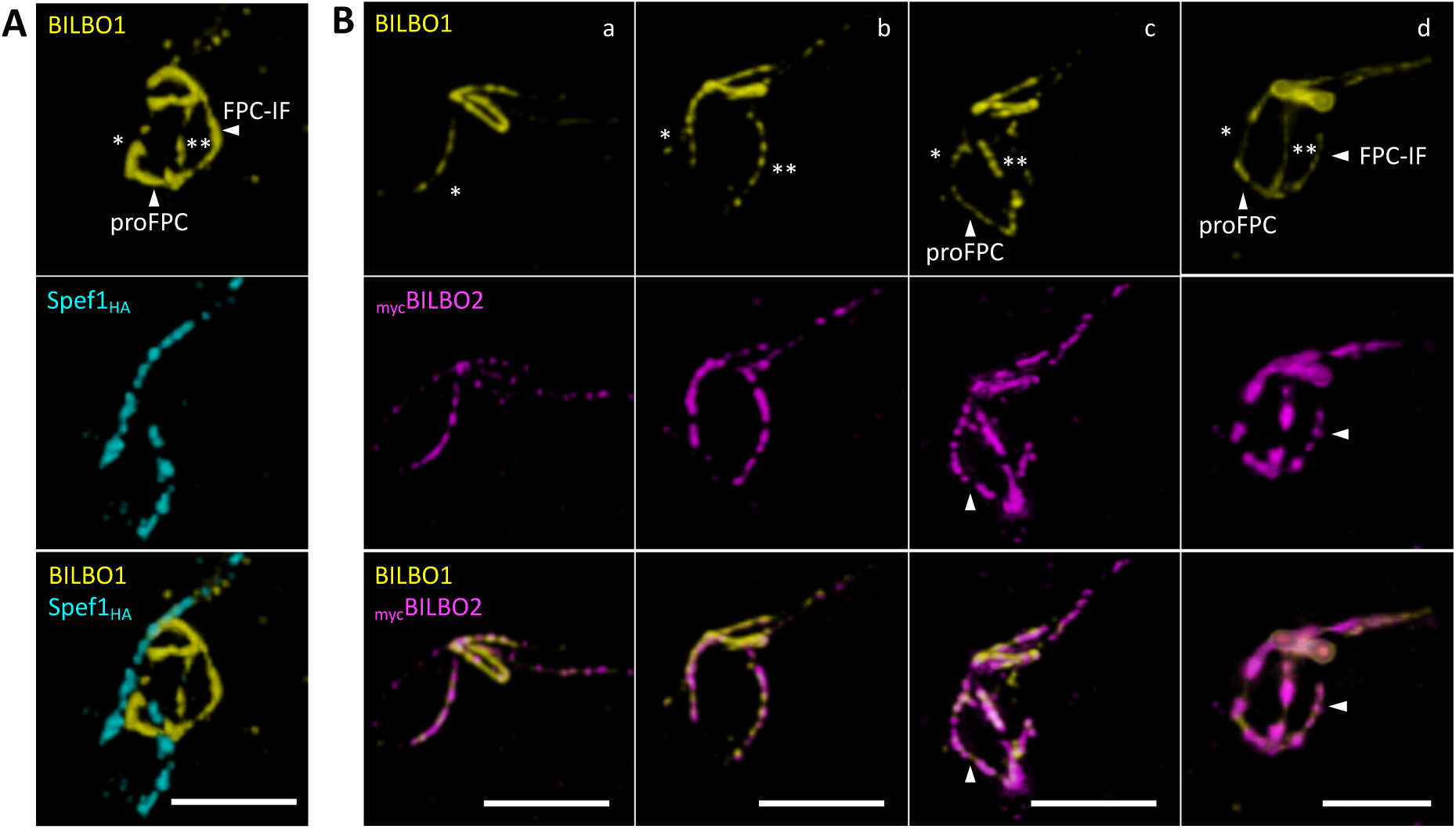
U-ExM immunofluorescence images showing that the proFPC and FPC-IF both contain FPC-specific proteins. **A**. Co-labelling of BILBO1 (yellow) and Spef1 (cyan). **B**. Co-labelling of BILBO1 (yellow) and BILBO2 (magenta). Scale bars: 5 μm.

Further investigations revealed that BILBO2, a direct interacting partner of BILBO1 and a known FPC component (15), colocalizes with BILBO1 along both the new and old MtQ throughout the cell cycle (Figure 5Ba, b). Notably, BILBO2 was also detected in the proFPC and the FPC-IF, reinforcing that these structures consist of specific, highly organized molecular components rather than transient, amorphous assemblies (Figure 5Bc, d). These findings underscore the specialized nature of the proFPC and FPC-IF and their important role in the cytoskeletal organization of *T. brucei*.

### BILBO1, MORN1, and BILBO2 localize to a cytoskeletal structure adjacent to the MtQ

To better understand the spatial organization of Spef1 relative to the MtQ, we performed co-immunolabeling of tubulin and Spef1, followed by fluorescence intensity profiling (Figure 6A). The resulting Spef1 intensity profile revealed two distinct peaks flanking the central tubulin peak, suggesting that Spef1 preferentially localizes on specific microtubules within the MtQ.

**Figure 6.**
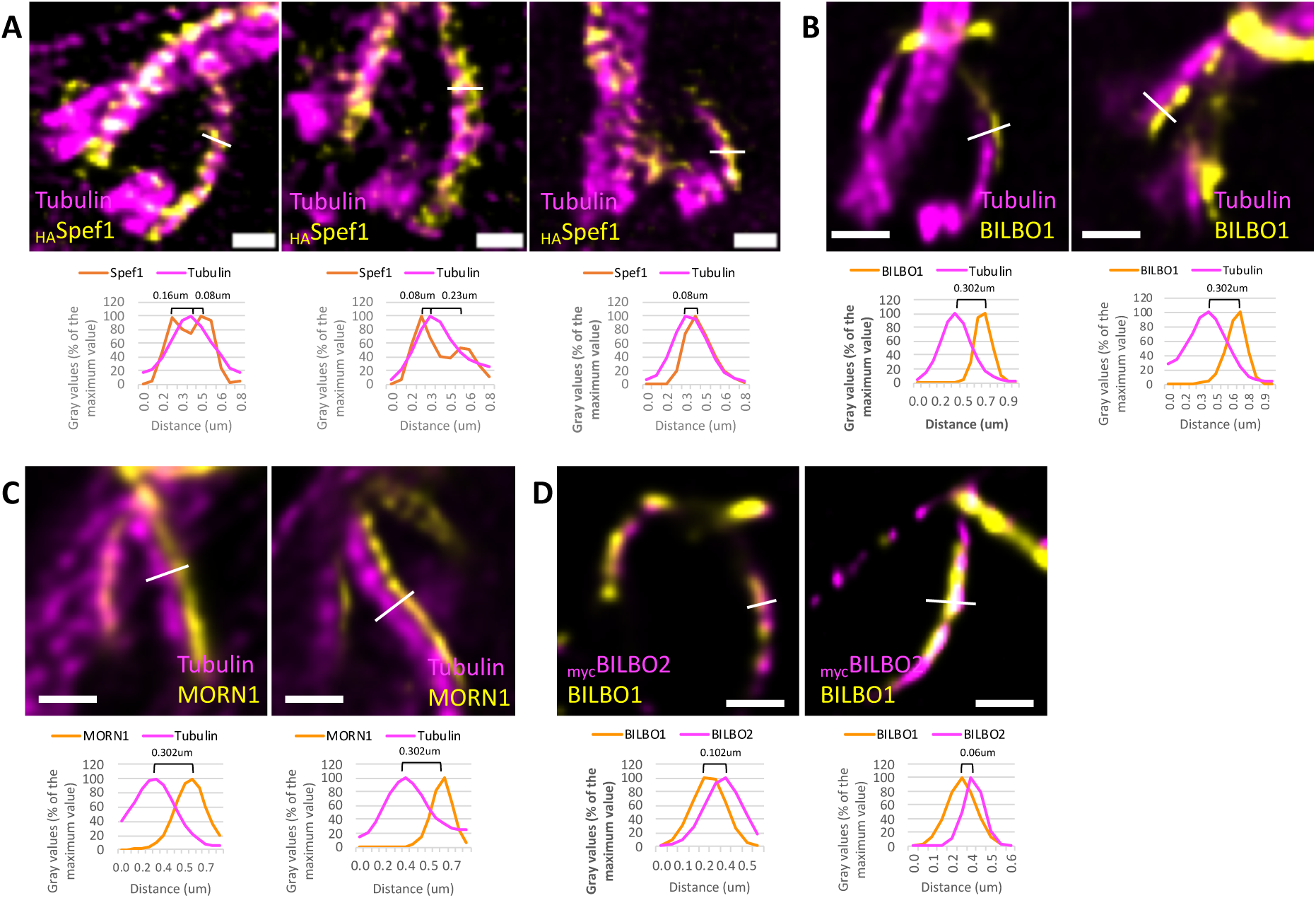
**A** U-ExM immunofluorescence images and intensity peaks measurements showing the spatial relationship between the MtQ (tubulin, magenta) and Spef1 labelling. **B.** Images and intensity peaks measurements comparing the MtQ (tubulin, magenta) and the BILBO1-positive filament. **C.** Images and intensity peaks measurements of the MtQ (tubulin, magenta) and the MORN1-positive filament. **D.** Images and intensity peaks measurements of the BILBO1-positive filament (yellow) and BILBO2 (magenta). Scale bars: 1 μm.

Measurements of the peak pixel intensities of tubulin and Spef1 showed a separation of 0.08 to 0.16 µm. Considering an expansion factor of 4.2× and the 25 nm diameter of a microtubule, this corresponds to an *in vivo* distance of approximately 20–40 nm. These results indicate that Spef1 is enriched along the two distal microtubules of the MtQ rather than being uniformly distributed across all four. In contrast, the distance between the peak pixel intensities of tubulin (MtQ) and either BILBO1 or MORN1 (Figure 6B, C), was approximately 0.3 µm, corresponding to roughly 70 nm *in vivo*. This suggests that BILBO1 and MORN1 are not directly localized on MtQ microtubules. Further co-labelling experiments with BILBO1 and BILBO2 revealed a smaller inter-peak distance of 0.06 to 0.12 µm (15–30 nm *in vivo*), suggesting that these proteins co-localize within a distinct structure adjacent to the MtQ.

These findings suggest that BILBO1, MORN1, and BILBO2 positioned between the basal body and the FPC, localize to a cytoskeletal structure or protein complex associated with the MtQ, but are not directly integrated into it. Alternatively, BILBO1 and MORN1 could be confined to a single microtubule within the MtQ, but this hypothesis is inconsistent with the observed Spef1 localization pattern and inter-protein distance measurements. Therefore, the data support the existence of a specialized cytoskeletal structure adjacent to the MtQ that houses these proteins.

### BILBO1 knockdown disrupts MtQ orientation but not its assembly

In BILBO1 RNAi-induced knockdown cells, key phenotypes include partial detachment of the nFg at the distal end, the failure to assemble a new FPC and FP, and disorganization of the HC (6). Initial electron microscopy studies of thin sections from BILBO1 RNAi cells did not detect the presence of a newly formed MtQ associated with the nFg and basal body (6,16). To further investigate whether the formation of the nMtQ depends on BILBO1, we re-examined the BILBO1 RNAi phenotype in expanded samples, using anti-tubulin and anti-MORN1 antibodies. In wild-type (WT) cells, each flagellum is associated with an MtQ and an HC (Figure 7Aa). However, in BILBO1 RNAi-induced cells, MORN1 and tubulin labelling revealed that the detached new flagellum remains linked to a pro-basal body and a tubulin-positive structure, likely corresponding to the nMtQ (Figure 7B). This finding suggests that MtQ formation can proceed independently of BILBO1 and the FPC.

**Figure 7.**
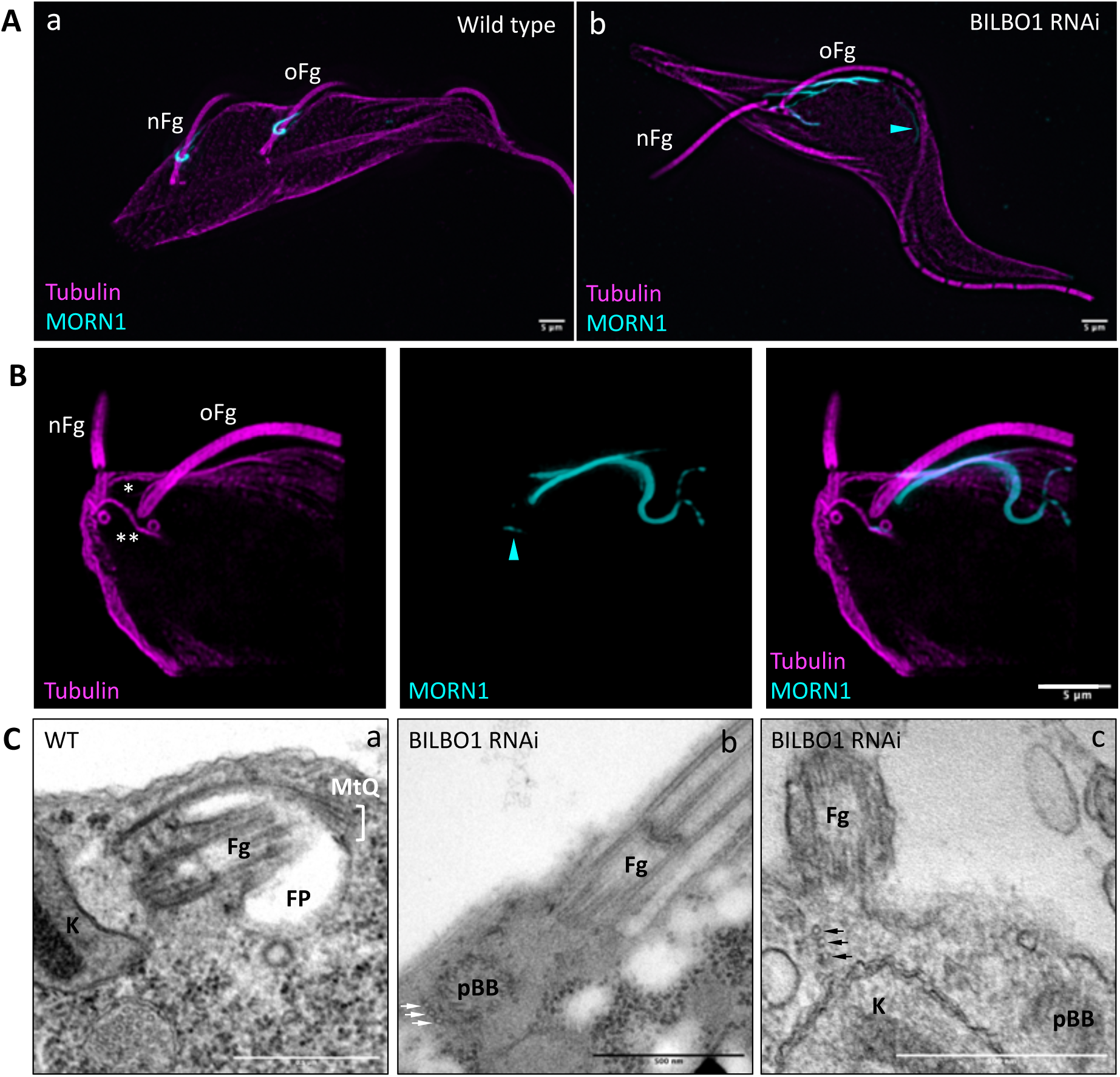
MORN1 (yellow) and tubulin (magenta) labelling on WT cells and BILBO1 RNAi-induced cells. **A.** Epifluorescence acquisition images and Z-project Max intensity representations. (a) Expanded detergent-extracted WT cells showing the hook shape of MORN1 labelling on both the old (oFg) and the new flagellum (nFg) (z-project Max intensity). (b) BILBO1 RNAi-induced cell displaying disrupted MORN1 localization and detachment of the new flagellum. Scale bars: 5 µm. **B.** Confocal acquisition of a detergent-extracted BILBO1 RNAi-induced cell exhibiting an advanced phenotype, where the new flagellum is detached along the cell’s length. A newly formed MtQ (**) is observed near the BB of the new detached flagellum but has an atypical orientation. MORN1 labelling is also abnormal at the old flagellum, with weak labelling along the nMtQ (arrowhead). Scale bar: 5 µm. **C.** Electron micrographs of thin sections of WT cells (a) and of BILBO1-RNAi induced cells (b, c). Arrows illustrate sets of intracellular microtubules resembling the MtQ but with three microtubules instead of four. Fg: flagellum; K: kinetoplast; proBB: pro-Basal body; FP: flagellar pocket. Scale bars: 500 nm.

Interestingly, in BILBO1 RNAi-induced cells (*e.g.,* Figure 7B), the nMtQ and its associated MORN1-positive component exhibited a misoriented trajectory, deviating from the typical alignment along the FP and the flagellum attachment zone. These observations indicate that while MtQ biogenesis is independent of the nFPC, its correct spatial orientation relies on the FPC. This dependency also extends to the HC structure. Moreover, since no new FPC was formed, it is possible that the proFPC or FPC-IF were either incomplete or absent, suggesting an indirect requirement for BILBO1 in their formation.

Longitudinal electron microscopy sections of WT cells, taken from the proximal region of the flagellum (where the central pair is visible), rarely reveal transverse sections of the MtQ. However, in one example (Figure 7Ca), the MtQ appears as an electron-dense region adjacent to the FP membrane. In contrast, in BILBO1 RNAi cells, the absence of an FP correlated with the mis-localization or partial displacement of the MtQ (Figure 7C, b, c). These findings further support the idea that BILBO1 is essential for the correct positioning and orientation of the MtQ even though its initial assembly does not require BILBO1.

## Discussion

In this study, we employed ultrastructure expansion microscopy (U-ExM) to investigate the biogenesis of the flagellar pocket collar (FPC) in *Trypanosoma brucei*, revealing previously unresolved details of FPC and HC formation. When combined with NHS protein labelling, U-ExM enabled the imaging of novel transient structures involved in FPC/HC assembly and provided the essential cellular context to complement antibody-based protein localization. The high-resolution capabilities of U-ExM offered new insights into the dynamic processes underlying FPC formation. Using these techniques, we uncovered critical interdependencies between the FPC, hook complex (HC), and microtubule quartet (MtQ). These findings redefine the molecular composition, functional roles, and dynamic processes of FPC and FP biogenesis.

### FPC biogenesis occurs de novo

While the spatial coordination of events leading to flagellar pocket formation and division was elegantly illustrated in previous work (28), the specific mechanisms of FPC biogenesis and function remained elusive. As schematized in Figure 8, our findings refine their model, demonstrating that FPC biogenesis occurs *de novo*, initiating at the pro-basal body (proBB) beneath the FP membrane. A nascent MtQ (nMtQ) extends along the proBB. Notably, previous work demonstrated that the nBB and nMtQ are decorated with BILBO1, even though the transition fibre protein TFK1 is not yet detected at the proBB (14,29). This suggests that nMtQ formation precedes proBB docking, or both events occur concomitantly. As the nMtQ extends and wraps around the FP, it unites with the old FPC. This is followed by the *de novo* formation of a BILBO1-MORN1-BILBO2-positive filament (the FPC-IF), which connects the proFPC to the oFPC at the anterior side of the FP. Just before or during pro-basal/BB body docking at the FP membrane, the old MtQ (oMtQ) also becomes decorated with these three proteins. Both MtQs are then linked *via* the semi-circular proFPC structure, which originates near the nBB transition fibres beneath the FP and serves as the foundation for the new FPC.

**Figure 8.**
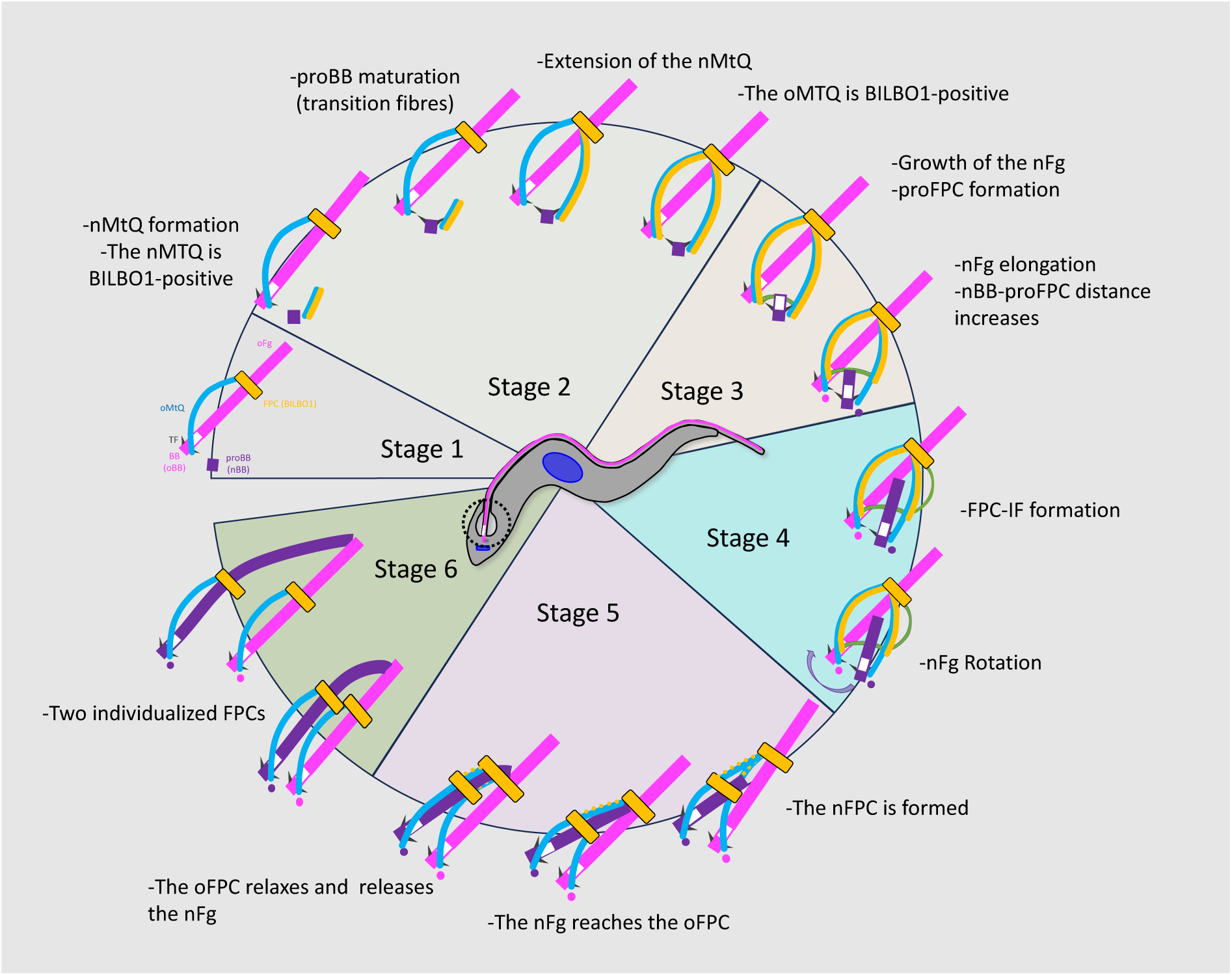
Schematic representation of FPC biogenesis stages. Stage 1, G1 cells with a single FPC and MtQ. **Stage 2**, The nMtQ begins to form, followed by proBB maturation. The nMtQ is decorated with BILBO1, subsequently, the oMtQ also acquires BILBO1. **Stage 3**, The nFg elongates alongside proFPC formation. **Stage 4**, As the nFg reaches the oFg - likely connecting *via* the flagella connector - the FPC-IF structure forms, followed by nFg-BB rotation. **Stage 5**, the nFPC fully develops, and the nFg exits the oFPC. In **Stage 6**, the two FPCs and flagella segregate in preparation for cytokinesis.

As the new flagellum (nFg) elongates within the FP, the distance between its new basal body and the proFPC increases. Additionally, the basal body undergoes rotary movement towards a more posterior cell position. While the precise mechanism underlying inter-basal body movements remains elusive, our data suggest a dynamic interplay between flagellum elongation and FPC remodelling. Several possibilities exist: (i) the proFPC moves along the flagellum, contributing to ridge and new FP formation, (ii) the proFPC remains stationary while the nFg elongates, or (iii) a combination of both, where the proFPC initially stays in place but later moves along the flagellum as it extends. Our data support the third hypothesis, in which the proFPC remains static at first but later migrates as the flagellum elongates. When the growing nFg approaches the boundary of the oFPC, the FPC-IF is formed between the proFPC and oFPC, linking the two structures. As the new flagellum rotates posteriorly, the proFPC closes into a corkscrew-like structure to form the mature nFPC. The FPC-IF then rapidly disappears, though whether this occurs *via* coordinated disassembly or degradation remains unknown. This disappearance marks the final separation of the two FPC structures.

Importantly, we show that the modification and relaxation of the oFPC into an open circular structure allows the new flagellum to exit the oFPC, while its distal tip remains attached to the oFg *via* the flagella connector. The oFPC must then restructure into its original compact, corkscrew-like conformation. This process requires precise coordination of key proteins, though the exact mechanism remains unknown. These findings align with research from the Gull lab, supporting the hypothesis that FPC biogenesis occurs *de novo* (30).

### MtQ biogenesis is independent of the FPC and required for flagellum attachment

We observed that nMtQ formation and extension occur before significant flagellum elongation, suggesting that MtQ biogenesis precedes flagellum growth. This is in agreement with previous findings on early cytoskeletal formation in trypanosomes (20). In BILBO1 RNAi knockdown cells, proFPC and FPC-IF formation is disrupted, yet the nMtQ still forms near the pro-basal body. This suggests that MtQ biogenesis is independent of the FPC, HC, and FP but relies on an alternative, unidentified mechanism. Furthermore, the absence of the flagellum attachment zone (FAZ) in BILBO1 RNAi cells (6), which leads to flagellum detachment along the cell length, suggests that the MtQ plays a key role in FAZ formation and positioning.

While FPC and HC proteins are associated with the MtQ, proFPC, and FPC-IF, no intraflagellar transport proteins have been identified at these sites. This suggests that if transport along the MtQ occurs, it does not use the classical IFT system, or that no transport is needed, with proteins instead recruited directly from the cytoplasm. However, several kinesins have been identified and localized in the proximal part of the MtQ and may facilitate the trafficking of FPC and HC components along the MtQ (14,24,31).

### Nature of BILBO1-positive structures and insights into HC and FP formation

Our study reveals that BILBO1 forms or contributes to a polymer running parallel to the MtQs and associating with the proFPC and FPC-IF. The polymerization properties of BILBO1, demonstrated both *in vivo* and *in vitro* (12,13), along with its interaction with proteins such as BILBO2 (14–17), suggest that BILBO1 may be a key candidate for polymer formation along the MtQs and for *de novo* structures such as proFPC and FPC-IF.

Our findings also suggest that the BILBO1-MORN1-BILBO2-positive structure at the nMtQ corresponds to the previously described “tendril” (10), which facilitates FPC biogenesis, parallels the nMtQ, and contains a dynamic pool of MORN1. We propose that BILBO1 provides a molecular skeleton for protein recruitment during FPC biogenesis (14,15,17,12). In BILBO1 RNAi knockdown cells, the absence of BILBO1 at the proFPC stage disrupts proper nMtQ orientation and prevents recruitment of key proteins, such as FAZ and interstitial zone layer proteins (32). Additionally, HC formation is impaired, as MORN1 and BHALIN (an HC protein and BILBO1 partner) become mis-localized (17,33). These findings highlight BILBO1 as a central organizer of multiple FP-associated structures. Our model aligns with previous studies (34), showing that FP formation precedes flagellum separation during proventricular stage development. However, key questions remain regarding the forces driving ridge invagination.

### Unresolved questions in FPC and HC formation

The mechanisms driving the initiation of the semicircle proFPC structure and the processes that generate the template linking the new and old MtQs remain unclear. BILBO1, which interacts with multiple proteins, may recruit factors essential for membrane docking and FPC-IF formation. Additionally, it may associate with motor proteins that drive proFPC movement along the axoneme, contributing to new FP formation. Notably, BILBO1 binds to TbKINX1B, a motor protein found on the FPC, BB and MtQ (14). While TbKINX1B knockdown did not prevent FPC biogenesis in cultured procyclic cells, it caused FPC reorientation to the anterior side of the nucleus, suggesting a role in FPC positioning. In bloodstream forms, TbKINX1B is essential, and its knockdown resulted in formation of enlarged flagellar pockets, further supporting an FP/FPC-related function. The exact trigger for proFPC movement remains unknown, but this sliding mechanism likely involves motor proteins that position the maturing FPC closer to the cell surface. This repositioning may facilitate ridge formation, a key step in FP formation.

### Conclusion

This study provides key insights into the biogenesis of the key structures FPC, MtQ, and flagellum in trypanosomes. While MtQ formation occurs independently of BILBO1, its proper orientation depends on an intact FPC, underscoring the intricate interplay between these structures. Further investigation into the molecular mechanisms governing MtQ orientation and the role of the FPC-IF in FPC formation will enhance our understanding of FP and FPC biogenesis.

## Materials and methods

### Cell lines, culture, transfection

Procyclic (PCF) cell lines SmOxP427 and SmOxP927 co-expressing the T7 RNA polymerase and tetracycline repressor (35) were grown at 27°C in SDM-79 medium supplemented with 2-2.5 mg/mL hemin, 26 µM sodium bicarbonate, 10% (v/v) complement-deactivated FBS, and 1 µg/mL puromycin. For transfection, cells were grown to 5×10^6^–1.0×10^7^ cells.mL^−1^, then 3×10^7^ cells were electroporated using transfection buffer as previously described (Wirtz et al., 1999; Schumann Burkard et al., 2011) with 10 µg of PCR product using the program X-001 of the Nucleofector®II, AMAXA apparatus (Biosystems). After transfection, clones were obtained by serial dilution and maintained in logarithmic phase growth. Selection antibiotics were added to the media according to the transfected product (neomycin 10 µg/mL, hygromycin 25 µg/mL, phleomycin 5 µg/mL).

The BILBO1 stem-loop RNAi vector p3960SL was previously described in (6). BILBO1 RNAi was induced for 48H with 10 µg/mL tetracycline. Endogenous 10×Myc-Nter tagged BILBO2 cell line was described in (15). Spef1 (10xHA tag at its N-terminus or C-terminus) and centrin arm associated protein 1, CAAP1 (10xmyc at its N-terminus) were endo-tagged using the pPOTv7 vector as described in (36).

### Immunofluorescence

#### Wide-field and epifluorescence microscopy

Cells were harvested, washed, and processed for immunolabelling on methanol-fixed detergent-extracted cells (cytoskeleton, CSK) as described in (16). The antibodies used and their dilutions are listed in Table S1. Images were acquired on a Zeiss Imager Z1 microscope, using a Photometrics Coolsnap HQ2 camera, with a 100× Zeiss objective (NA 1.4) using Metamorph software (Molecular Devices), and processed with ImageJ (NIH). The U-ExM wide-field images were deblurred with Metamorph Deconvolution 2D.

#### Ultrastructure-Expansion microscopy

##### U-ExM on detergent-extracted cytoskeletons

The U-ExM protocol has been adapted from (37). Briefly, 4×10^6^ mid-log phase cells were washed in PBS and loaded onto poly-L-lysine 0.1% solution -coated 12 mm coverslips in 24-well plates and cells left to adhere for 5– 10 min. Cells were extracted for a few seconds with PBS NP40 (Igepal) 0.25%. Cells were then covered with 1 ml of activation solution (0.7% formaldehyde, 1% Acrylamide in PBS) for 3.5 h at 37°C. For the gelation step, the coverslips were gently deposited on top of a 35 µl drop of MS solution (23% sodium acrylate, 10% acrylamide, 0.1% bisacrylamide 0.5% TEMED and 0.5% ammonium persulfate in PBS) for 5 min then transferred at 37°C and incubated for 1 h in a moist chamber. The coverslips were then transferred to a 6-well plate in 1 ml of denaturation solution (200 mM SDS, 200 mM sodium chloride, 50 mM Tris-HCl pH 9.0) with agitation at RT for 15-30 min to detach the gel from the coverslip. The gel was then moved into a 1.5 ml Eppendorf centrifuge tube filled with denaturation solution and incubated at 95°C for 90 min. Gels were expanded in 100mL of deionized water (3 × 30 min) then incubated 3 × 10 min in 100mL of PBS. Small pieces of the gels were processed for immunolabelling as follows. The gels were preincubated in blocking solution (PBS, 1-2% BSA and 0.2% Tween20 or PBS only) for 30 min at 37°C, and then with the primary antibodies (see table S1 for the reference and dilutions) overnight at 37°C with shaking. After three washes in 1 mL blocking solution, the gels were incubated with the secondary antibodies diluted in blocking solution for 4.5 h in the dark at 37°C with slow agitation. After three washes in blocking solution, gels were expanded in 100 mL of deionized water (3 × 30 min). An expansion factor was determined using the ratio between the size of the coverslip (12 mm) and the size of the gels after the first expansion and was 4.2-fold.

##### U-ExM on whole cells

The U-ExM protocol on whole cells was performed as previously described in (21). Briefly, 2×10^6^ cells were washed 2x in PBS, fixed with 4% formaldehyde and 4% acrylamide in PBS, seeded onto 12 mm round coverslips washed in NaOH, and incubated overnight. Next day, the coverslips were gently washed in PBS and inverted cell side down onto a 50 µL drop of polymerizing monomer solution (19% sodium acrylate, 10% acrylamide, 0.1% N,N’-Methylenebisacrylamide, 0.5% TEMED, 0.5% ammonium persulfate in PBS). The gels were allowed to settle for 5 min at RT, then incubated at 37°C for 30 min. The coverslips were then carefully removed from the gels by slight expansion of the gels in dH_2_O, and the gels were immediately transferred into denaturation buffer and incubated at 95°C for 1 h. After denaturation, gels were expanded by washing in dH_2_O 3 × 20 min and subsequently shrunk in PBS for 30 min. For antibody staining, 1/4 of the gel was cut with a razor and incubated in primary antibody diluted in blocking buffer (2% BSA, 0.02% sodium azide in PBS) 6 h to overnight. The unbound primary antibody was removed by washing in PBS 3 × 20 min and the gels were subsequently incubated in secondary antibody diluted in blocking buffer 6 h to overnight. The unbound secondary antibody was removed by washed 3 × 20 min in PBS and the gels were incubated 90 min in 20 µg/mL NHS ester-ATTO 594 in PBS. Finally, the gels were washed 3 × 20 min in PBS and expanded 2 × 30 min in dH_2_O prior to imaging.

#### Confocal microscopy

For imaging expanded cytoskeletons, we used a Leica SP8 on an upright stand microscope DM6000 (Leica Microsystems, Mannheim, Germany), with a HCX Plan Apo CS 63X oil NA 1.40 objective. The lasers used were Diode 405 nm, OPSL 488, OPL 552 nm and Diode 638. The system was equipped with a conventional scanner (10Hz to 2800 Hz) and four internal detectors (2 conventional PMT and 2 hybrid detectors). The images were deconvolved in Huygens Pro and processed with ImageJ and Fiji software (38,39). The quantification in figure 2G was obtained as follows. Eighteen slices of U-ExM confocal images were flattened with Fiji (Z-project SUM), and two areas were manually defined: a close area around the old FPC (excluding the BILBO1 labelling distal to the FPC), and the area from the old FPC to the basal bodies and including the old FPC and the MtQs. Figure 2G indicates the integrated fluorescence of BILBO1 at the old FPC as a percentage of the total fluorescence in the total area.

For imaging expanded whole cells, a Leica DMi8 with TCS SP8 confocal head (Leica Microsystems, Mannheim, Germany) equipped with the HC Plan APO CS2 63x oil NA 1.40 objective, a 20 mW 488 nm and a 20 mW 552 nm solid-state lasers, a standard scanner (1-1800 Hz line frequency), and 5 detectors (2 × PMT and 3 × HyD), and a Leica DMi8 with STELLARIS 8 confocal head, with HC Plan APO 86x water immersion NA 1.20 objective, wide-range white light laser with pulse picker (WLL PP), conventional linear scanner (1-2600 Hz line frequency) and 5 supersensitive hybrid detectors were used. The images were deconvolved using Huygens professional. To visualize the stages of FPC biogenesis, the data were segmented in Bitplane Imaris software by manual segmentation combined with intensity thresholding.

#### Electron microscopy

A total of 50 ml of mid-log phase WT or 48-h BILBO1 RNAi-induced cells were harvested by centrifugation at 1,000 × g for 15 min. Block preparation and EM protocol was performed as in (40).

## Abbreviations

BB: basal body
FP: flagellar pocket
FPC: flagellar pocket collar
FPC-IF: flagellar pocket collar interconnecting fibre
HC: hook complex
MtQ: microtubule quartet
nBB: new basal body
nFg: new flagellum
nFp: new flagellar pocket
nHC: new hook complex
nMtQ: new microtubule quartet
oBB: old basal body
oFg: old flagellum
oHC: old hook complex
oMtQ: old microtubule quartet
proBB: pro-basal body
proFPC: pro-flagellar pocket collar
U-ExM: ultra-structure expansion microscopy

## Acknowledgments

We thank the ProParacyto group members for stimulating discussions, and Eloïse Bertiaux for critical reading of the manuscript. We thank B. Morriswood for the anti-MORN1 antibody, K. Ersfeld for the 9E10 anti-myc antibody, and S. Dean for the pPOTv7 vectors. We thank Eloïse Bertiaux and Martin Bablon for their help to design the scheme in figure 8. We also acknowledge the Bordeaux Imaging Centre, a service unit of the CNRS-INSERM and Bordeaux University, a member of the national infrastructure France BioImaging supported by the French National Research Agency (ANR-10-INBS-04). The help of Christel Poujol, Magali Mondin and Mónica Fernández Monreal is acknowledged. We also acknowledge the Light Microscopy Core Facility, IMG, Prague, Czech Republic, supported by MEYS – LM2023050 and RVO – 68378050-KAV-NPUI, for their support with the confocal imaging of expanded whole cells presented herein.

---

**Table S1.**

| Antibody | Provider / Reference | Dilution Non-expanded | Dilution U-ExM | Figure |
| --- | --- | --- | --- | --- |
| **Primary antibodies** |  |  |  |  |
| MORN1 rabbit polyclonal | (41) | 1:4,000 | 1:2,000 | All MORN1 figures |
| BILBO1 rabbit polyclonal (anti aa 1-110) | (12) | 1:2,000 | 1:1,000 | All BILBO1 figures except Figure 3 |
| BILBO1 rabbit polyclonal | (10) |  | 1:500 | Figure 3 |
| Rabbit anti-HA | GeneTex GTX115044 |  | 1:500 | Figure 1, Figure 6 |
| Mouse anti-tubulin | DM1A, Sigma T9026 |  | 1:100 | Figure 1, Figure 6 |
| Mouse anti-myc | Ozyme 2276S |  | 1:500 | Figure 6, 7 |
| Mouse anti-myc 9E10 | DSHB |  | 1:250 | Figure S1 |
| Mouse anti-HA | Santa cruz 7392 |  | 1:200 | Figure 7 |
| Guinea pig anti-tubulin | AA345-GP<br>Geneva Antibody Facility |  | 1:500 | Figure 2, 6, 8 |
| **Secondary antibodies** |  |  |  |  |
| anti-mouse A488 | Molecular Probes A11001 |  | 1:200 | Figure 1, Figure 6 |
| anti-mouse A647 | ThermoFischer A21235 |  | 1:500 | Figure 6, 7 |
| anti-mouse A546 | ThermoFischer A11030 |  | 1:500 | Figure 7 |
| anti-rabbit A594 | ThermoFischer A11012 |  | 1:500 | Figure 1, Figure 6 |
| anti-rabbit A488 | ThermoFischer A11008 |  | 1:500 | Figure 2, 4, 6, 7 |
| anti-rabbit A647 | ThermoFischer A21244 |  | 1:500 | Figure 8 |
| anti-guinea pig A488 | ThermoFischer A11073 |  | 1:500 | Figure 8 |
| anti-guinea pig A647 | ThermoFischer A21450 |  | 1:500 | Figure 2, 4, 6 |
| **NHS Ester** |  |  |  |  |
| ATTO 594 NHS ester | Sigma 08741 |  | 20 µg/mL in PBS | Figure 3, 4 |

**Figure S1.**
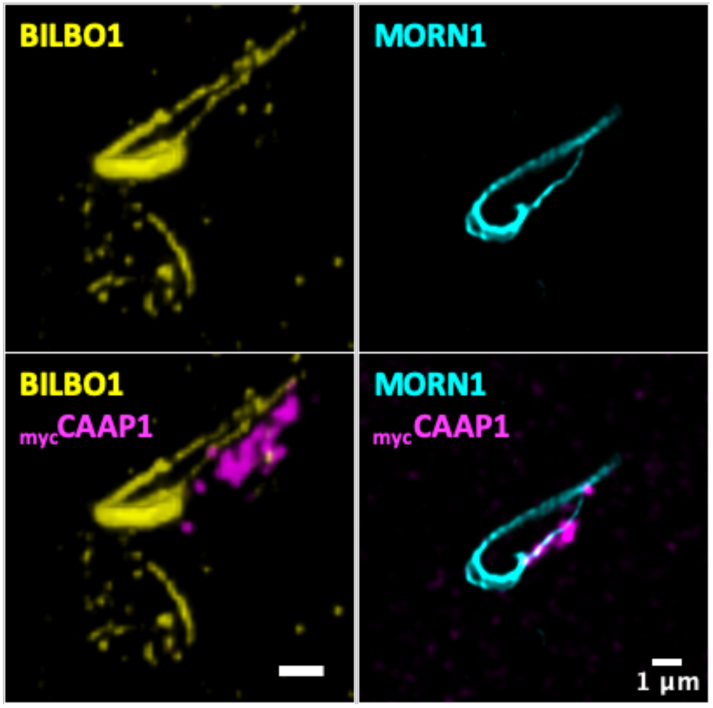
U-ExM immunofluorescence images showing that a sub-population of BILBO1 (yellow) and MORN1 (cyan) localize close to the centrin arm, which is indicated by _myc_CAAP1 labelling (magenta). Scale bars: 1 μm.

**Figure S2.**
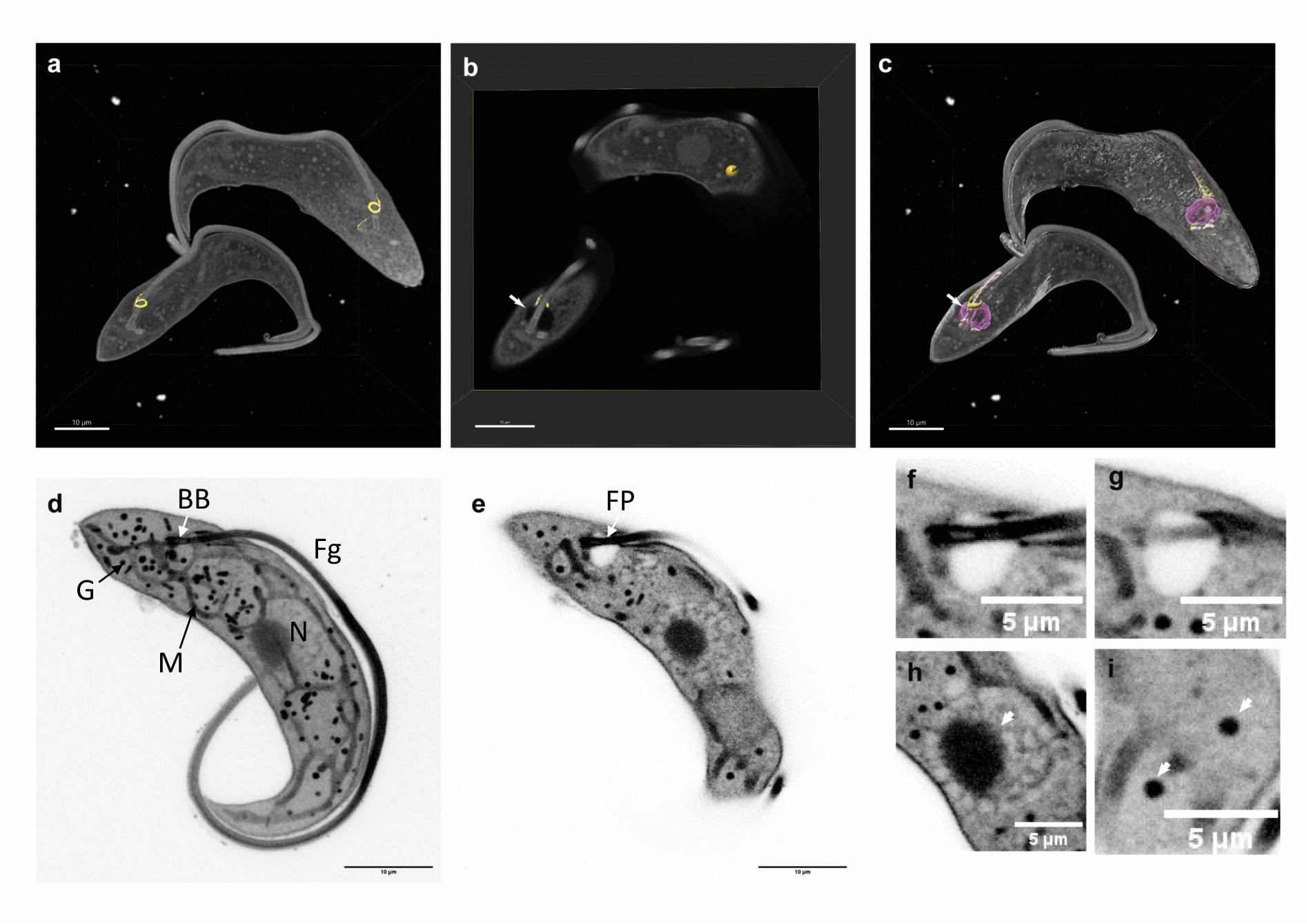
NHS ester labelling of whole expanded cells. **a** 3D rendering of two cells labelled with NHS ester (grey) and an antibody against BILBO1 (yellow). **b** A single Z-plane of the same data viewed through the orthoslicer. The arrow points to the flagellar pocket (FP), which is not visible in whole-cell rendering. **c** Segmentation of the data shown in (a) and (b). The cell surface (transparent grey) was segmented from the NHS ester signal by intensity thresholding. Microtubule-based structures at the flagellum base (solid grey) were segmented by manual masking combined with intensity thresholding. The FP (purple, arrow) was modelled by manual segmentation, and the BILBO1 signal (yellow) was segmented by intensity thresholding. **d** Maximum intensity projection of NHS ester labelling (inverted LUT). NHS ester labels organelles and structures within the cell, such as the nucleus (N), mitochondrion (M), glycosomes (G), the flagellum (Fg), and the basal bodies (BB). **e** A single Z-plane of the data in (d). Slicing through the volume reveals additional details, such as the shape of the flagellar pocket (FP). **f-i** Details of single Z-planes of the data represented in (d) and (e). **f,g** Details of the flagellar pocket. **h** Nucleus with visible nucleolus (arrow). **i** Detail of glycosomes (arrows). Scale bars: 10 μm in a-e, 5 μm in f-i.

